# Genomic analysis of allele-specific expression in the mouse liver

**DOI:** 10.1101/024588

**Authors:** Ashutosh K. Pandey, Robert W. Williams

**Author notes:** CORRESPONDING AUTHOR: Ashutosh K. Pandey 855 Monroe Avenue, Suite 512 Memphis, TN, 38163 Email addresses.

## Abstract

Genetic differences in gene expression contribute significantly to phenotypic diversity and differences in disease susceptibility. In fact, the great majority of causal variants highlighted by genome-wide association are in non-coding regions that modulate expression. In order to quantify the extent of allelic differences in expression, we analyzed liver transcriptomes of isogenic F1 hybrid mice. Allele-specific expression (ASE) effects are pervasive and are detected in over 50% of assayed genes. Genes with strong ASE do not differ from those with no ASE with respect to their length or promoter complexity. However, they have a higher density of sequence variants, higher functional redundancy, and lower evolutionary conservation compared to genes with no ASE. Fifty percent of genes with no ASE are categorized as house-keeping genes. In contrast, the high ASE set may be critical in phenotype canalization. There is significant overlap between genes that exhibit ASE and those that exhibit strong *cis* expression quantitative trait loci (*cis* eQTLs) identified using large genetic expression data sets. Eighty percent of genes with *cis* eQTLs also have strong ASE effects. Conversely, 40% of genes with ASE effects are associated with strong *cis* eQTLs. *Cis*-acting variation detected at the protein level is also detected at the transcript level, but the converse is not true. ASE is a highly sensitive and direct method to quantify *cis*-acting variation in gene expression and complements and extends classic *cis* eQTL analysis. ASE differences can be combined with coding variants to produce a key resource of functional variants for precision medicine and genome-to-phenome mapping.

## INTRODUCTION

Genetic variation contributes greatly to phenotypic diversity and differences in disease susceptibility by altering the structure and expression levels of proteins. The analysis of complex phenotypes in the pre-genomic era focused on coding variants, especially including nonsense, missense, and frameshift mutations. However genome-wide association studies conducted over the last decade have demonstrated that a great majority (>90%) of trait/disease-associated variants are located in non-coding regions. These non-coding variants primarily act by modulating gene expression, and they are the major cause of variation in susceptibility to complex diseases (Manolio *et al.* 2009; Maurano *et al.* 2012; Ward and Kellis 2012).

Sequence variants that affect gene expression can act in *cis* or in *trans*. *Cis*-acting variants represent first-order local control of gene expression that is specific to each individual haplotype. For example, sequence variants in transcription factor binding sites may affect expression of cognate genes on the same chromosome. Cis-acting variants are key to understanding heritable variation in disease risk, and serve as direct targets for diagnosis and treatment of diseases. Cis-acting variants can, of course, also have second-order distal or *trans* effects. A small subset of cis-modulated transcripts consists of master trans-regulators (for example, transcription factors, miRNAs) that control the abundance of large numbers of downstream target genes on both sets of chromosomes. Hence, genome-wide identification of cis-modulated transcripts serves as an important molecular resource for reverse genetics studies that focus on downstream consequences of altered gene expression.

Currently, two genome-wide approaches can be employed to identify cis-acting variation in expression. The first approach, known as expression quantitative trait locus (eQTL) mapping, performs classical genetic linkage analysis of expression usually for an entire transcriptome or proteome. This approach has been widely applied to study the effects of segregating variation on gene expression in yeast, mice, maize, and humans (Brem *et al.* 2002; Chesler *et al.* 2005; Crowley *et al.* 2015; Damerval *et al.* 1994; PEIRCE *et al.* 2003; SCHADT *et al.* 2003). The largest study to date is the ongoing Genotype-Tissue Expression (GTEx) project that is generating a comprehensive resource of *cis* eQTLs for multiple tissues in a large human cohort (KEEN and MOORE 2015; LONSDALE *et al.* 2013). The second approach, widely used in studies of model organisms, exploits rtPCR or RNA-seq to assay allele-specific expression (ASE) differences in isogenic heterozygous (F1) individuals (BELL *et al.* 2013; CIOBANU *et al.* 2010; MCMANUS *et al.* 2010; ROZOWSKY *et al.* 2011; SZABO and MANN 1995). RNA-seq can reliably distinguish mRNAs transcribed from the alternative alleles, and can be used to detect unequal production of the two alleles. A major advantage of isogenic F1 hybrids is that they provide a way to control for environmental and trans-acting influences. Both alleles are present within an identical environment and subjected to the same genetic background and regulatory networks. As a result, any expression differences between alleles in an isogenic F1 can be confidently attributed to genetic or epigenetic regulatory variant acting in *cis* (DEVEALE *et al.* 2012; LAGARRIGUE *et al.* 2013; WANG *et al.* 2008).

In this study we evaluate and compare the impact of cis-acting variation on expression in murine liver using both ASE and eQTL approaches. We exploit RNA-seq data from isogenic F1 hybrids and array data from a large set of recombinant inbred strains of mice—the BXD cohort, and generate a molecular resource for genome-wide reverse genetics that focuses on downstream consequences of altered gene expression (Carneiro *et al.* 2009; Li *et al.* 2010). We address the following questions:

- How do genes that exhibit ASE differ from those that do not?
- How do the two approaches highlighted above compare in terms of detecting effects of local polymorphisms on expression? More specifically, are cis-modulated transcripts identified by eQTL mapping also consistently detected by ASE analysis?
- How frequently do cis-acting variants that cause mRNA differences also cause differences in protein expression?

## RESULTS

### DBA/2J specific reference genome

We identified ∼4.46 m high confidence SNPs between C57BL/6J (*B*) and DBA/2J (*D*) genomes using 65-fold coverage next-generation sequencing data (Methods). We substituted these SNPs into the reference genome (GRCm38/mm10) to create a customized DBA/2J genome for RNA-seq read alignment (see below). A total of ∼1.7 m SNPs are located within coding genes based on RefSeq annotation (Table S1), including introns (95.59%), exons (2.30%), 3’ UTRs (1.80%) and 5’ UTRs (0.30%). These SNPs are distributed among 14,591 genes. SNPs in transcribed regions were used to discriminate between, and identify, the parental allelic origin (*B* vs *D*) of transcripts in isogenic F1 hybrids.

### Haplotype-aware alignment corrects for allelic bias in RNA-seq read alignment

We downloaded paired-end liver RNA-seq reads for six biological replicates of C57BL/6JxDBA/2J F1 females (Methods). We adopted a haplotype-aware alignment approach and aligned ∼350 m (∼175 m paired-end) reads against both the *B* and the customized *D* genomes (Methods, Table S2). We used a SNP-directed approach to identify the allelic origin (*B* or *D*) of reads that aligned over heterozygous SNPs in the F1 samples. Only uniquely aligned reads were assigned to parental alleles. Approximately 0.27 m SNPs within genes (a great majority within exons) had at least one read.

RNA-seq read alignment suffers from allelic bias that disfavors reads containing sequence variants relative to the reference genome (Degner *et al.* 2009; Satya *et al.* 2012). This bias generates lower read counts for non-reference alleles, and overestimates ASE differences. To evaluate bias, we examined allelic ratios—defined as the number of reads with the reference allele (*B*) divided by the total number of reads (*B*+*D*). In the absence of bias, this ratio will have a symmetrical distribution and a mean of 0.5. For each of the F1 samples, ratios were well balanced, with nearly equal numbers of SNPs with high *B* or high *D* expression (Table S3). Additionally, mean and median ratios were close to 0.5 (Table S3) indicating that the majority of SNPs exhibit small or undetectable ASE. We compared our results with a traditional approach involving alignment of reads against the reference genome and allowing for fewer mismatches (1 mismatch per 25 nt). This produced an artifactually high number of SNPs with high *B* expression (∼ 3,000 *B* vs ∼400 *D*, two-sided binomial *p* < 10^-323^, Fig. 1A) compared to the dual genome alignment (∼2,325 *B* vs ∼ 2,300 *D*, two-sided binomial *p* value = 0.724, Fig. 1B). The mean and median of allelic ratios using the standard approach were also skewed—0.69 (high *B*) and 0.68, respectively. This illustrates that the haplotype-aware alignment workflow is highly effective in reducing allelic bias.

**Figure 1.**
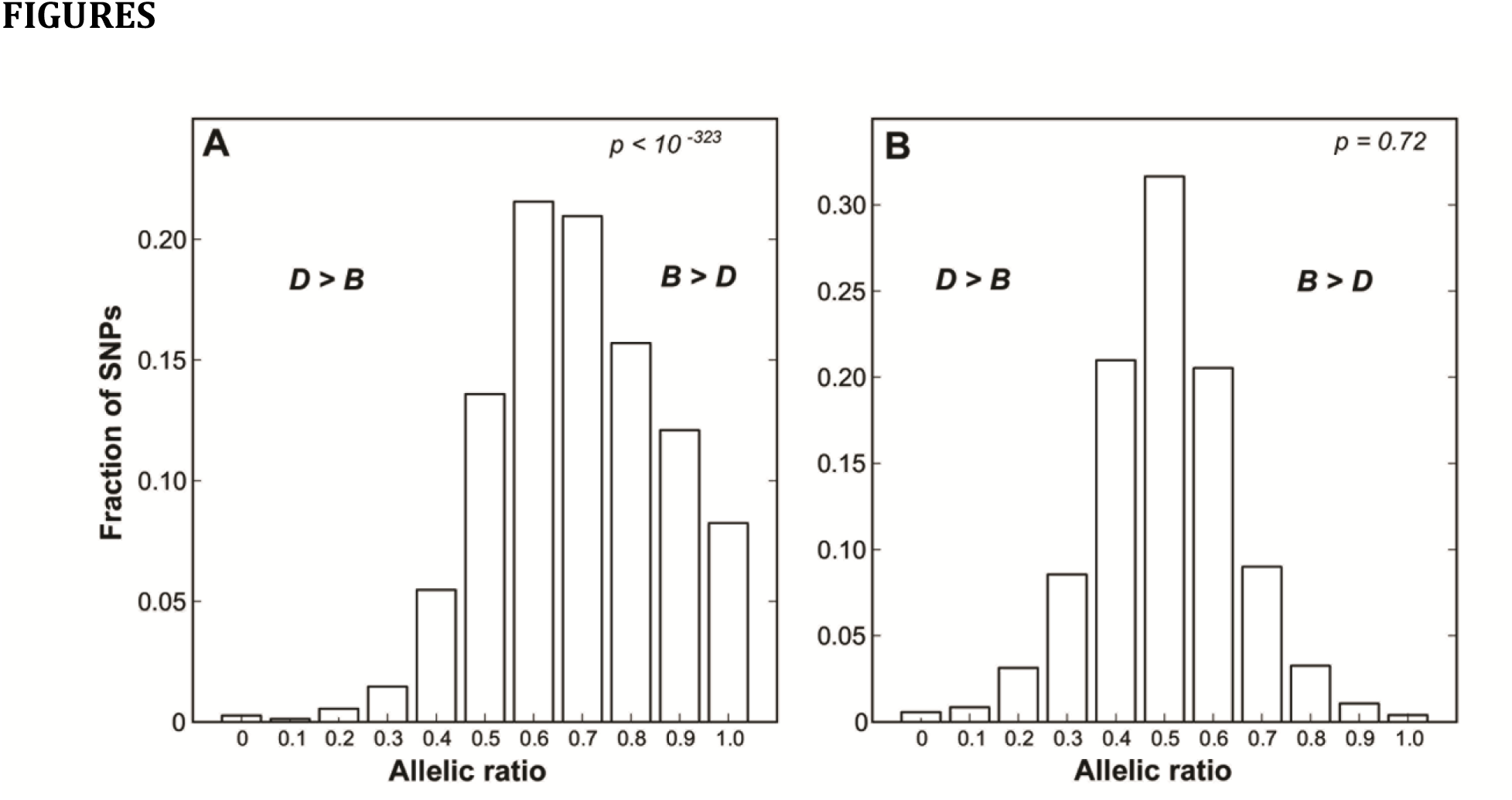
Comparison of the allelic bias in read alignment between traditional and haplotype-sensitive approach. Distribution of allelic ratios in (A) traditional and (B) haplotype-sensitive alignment.

### High correlation of allelic ratios across biological replicates

We calculated the correlation of allelic ratios with read depth ≥ 20 across all biological replicates. Allelic ratios were highly correlated with an average Pearson correlation of 0.70 ± 0.02 for all pairs of replicates (*n* = 15, Table S4, Fig. S1). We merged data from biological replicates, but to minimize variation across replicates, we discarded reads from replicate with highly discrepant ratio (Methods). More than 90% of SNPs had closely matched ratios across four or more replicates and were retained for further analyses. We also checked the concordance of the polarity of ASE measured by neighboring but independent SNPs (<75 nucleotides). In the great majority of cases SNPs within the same genomic feature (5’ UTR, exon, intron and 3’ UTR) were highly concordant. Only 136 (6%) of 2,234 genomic features contained SNPs with opposite ASE polarity and the great majority were in 3’ UTRs (*n* = 86). 3’ UTRs undergo extensive alternative processing (Hilgers *et al.* 2011; Miura *et al.* 2013), and SNPs with opposite ASE polarity probably represent alternative polyadenylation sites (Li *et al.* 2010). The high correlation of allelic ratios across the replicates and the high concordance in polarity again demonstrate the accuracy of the haplotype-aware alignment workflow. SNPs located within copy number variants, large insertions and deletions or in close vicinity (< 75 nucleotides) to an indel can generate inaccurate ASE estimates due to alignment artifacts. They were removed from further analysis. Additionally, to ensure independent sampling we considered only one SNP of a SNP pair when SNPs were separated by less than 75 nucleotides. Of 21,166 SNPs with read coverage ≥ 30, ∼25% (5,358) SNPs were removed for one of these reasons.

### ASE differences in liver are common

We tested the null hypothesis of equal abundance of transcripts representing *B* and *D* alleles in isogenic F1 hybrids using a Chi-square Goodness of fit test (FDR < 0.1, Methods). On average we used ∼650 reads per SNP to test for ASE. At a minimum threshold of 30 reads per SNP, we were able to test 15,808 SNPs in 3,589 genes (Table S5). We detected significant ASE in 5,298 SNPs from 1,905 genes (Table S6, Table S7 Fig. 2). Most of these SNPs are contained within coding exons (40%) and 3’ UTRs (40%) (Table S6). Seven percent are in introns and may represent unannotated exons or transcripts with unspliced or retained introns. We obtained comparable results when the minimum read threshold was increased to ≥ 60 (Table S6, 4,968 SNPs in 1,791 genes) and when the FDR threshold was decreased to 0.05 (4,774 significant SNPs in 1,482 genes). Fifty-two percent (2,773 *B* vs 2,525 *D*) of SNPs have higher expression from the *B* allele. There is no difference in the distribution of average ASE effect sizes between alleles (Fig. 3). Over half of the SNPs with significant ASE differ by less than two-fold; one-third differ 2–4 folds; and the remaining one-sixth differ more than 4-fold. We detected the known low *D* allele expression of aryl hydrocarbon receptor (*Ahr*)—a transcription factor that controls xenobiotic metabolizing enzymes (Lin *et al.* 2011). Similarly we also detected the known low *B* allele expression of alkaline phosphatase (*Alpl*)—a gene linked to hypophosphatasia (ANDREUX *et al.* 2012).

**Figure 2.**
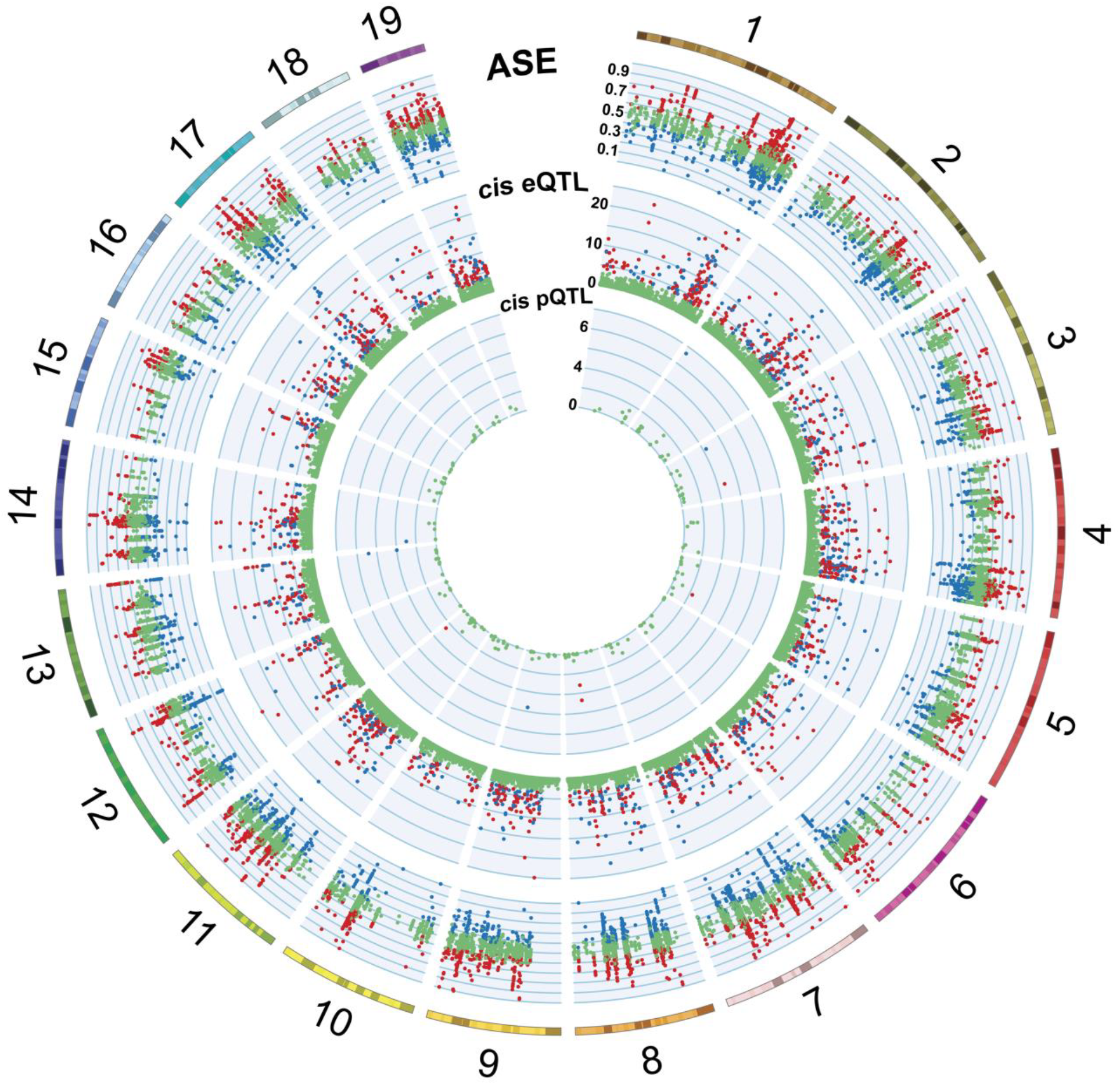
CIRCOS plot showing distribution of cis-modulated genes. The outermost circle represents chromosomes. Moving in, the second circle represents a scatter plot of ∼15,000 SNPs tested for ASE. The *Y*-axis represents the allelic ratio. SNPs with significant ASE are shown in red and blue, representing high expression of the *B* and the *D* allele respectively. Insignificant SNPs showing equal abundance of the *B* and the *D* allele are shown in green. These SNPs are located on or near the line representing an allelic ratio of 0.5. The third circle represents a scatter plot of ∼40,000 microarray probe sets tested for *cis* eQTLs. The *Y*-axis represents the LOD scores of probe sets measured at the nearest marker (*cis* LOD). *Cis* LOD (≥ 3) scores associated with high expression of the *B* and the *D* allele are shown in red and blue respectively. *Cis* LOD scores of less than 3 are shown in green. The innermost circle represents a scatter plot of ∼200 proteins tested for *cis* pQTLs. The *Y*-axis represents the LOD scores of proteins measured at the nearest marker (*cis* LOD). Chromosomes X and Y were excluded.

**Figure 3.**
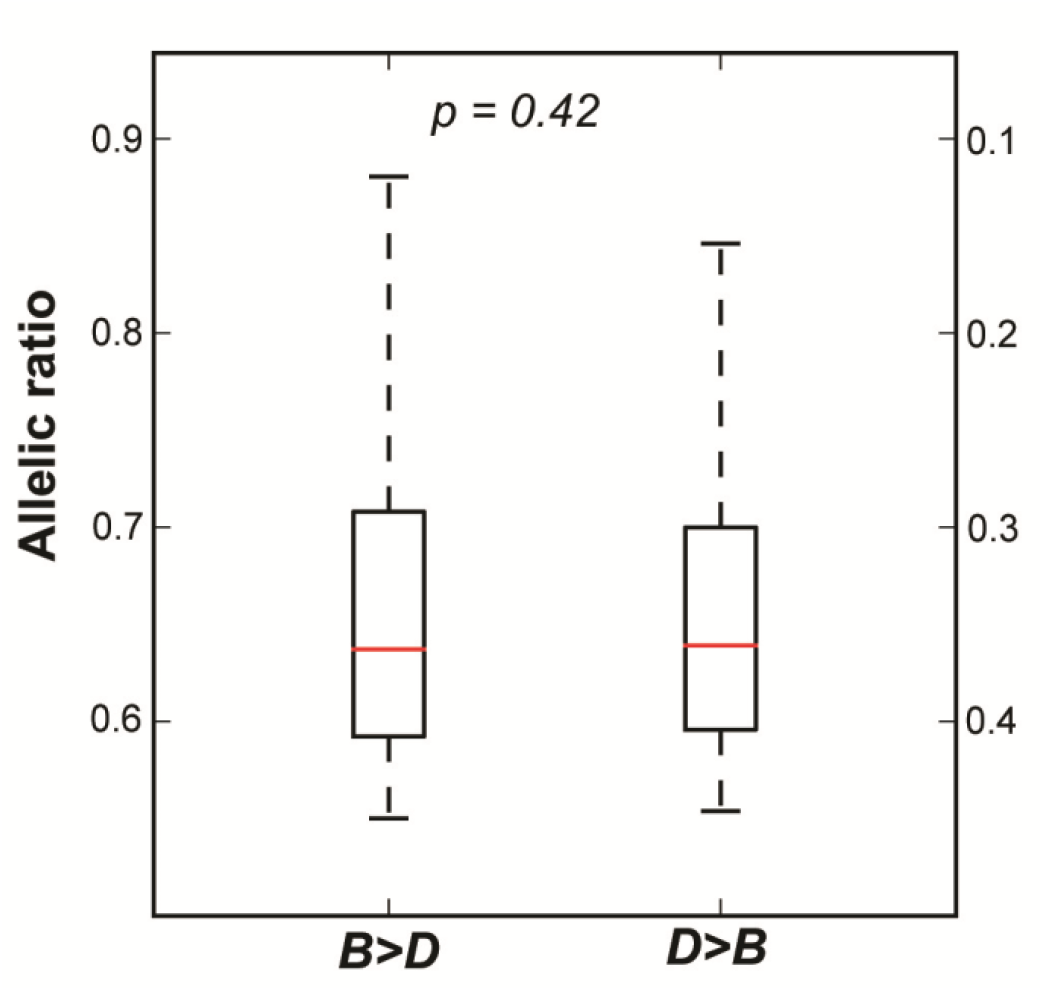
Distribution of allelic ratios. The left boxplot labelled as “*B*>*D*” represents SNPs with high expression of the *B* allele (left *Y*-axis). The right boxplot represents SNPs with high expression of the *D* allele (right *Y*-axis). The *Y*-axis represents allelic ratios. Outliers are not shown.

### Comparison between ASE and non-ASE genes

Genes with high or low levels of ASE may differ in length, complexity of promoters, sequence variant density, or evolutionary history. To explore these differences we selected a subset of 418 genes with very high ASE ratios (>1.5 fold) and a subset of 465 genes with low or no ASE. All genes in both groups were required to have at least two independent SNPs that supported their categorization. We also required all SNPs to have more than 100 supporting reads—roughly the top ten percentile. We defined each gene as the region between the transcription start site and 3’ UTR with 2 kb of flanking regions upstream and downstream. ASE genes do not differ from non-ASE genes in terms of total gene length or their 5’ or 3’ UTR length (Table 1). They also do not differ in numbers of protein-coding transcripts (isoforms) or numbers of exons per transcript (Table 1). However, ASE genes have a higher functional redundancy (number of paralogs) compared to non-ASE genes (1.5 fold, *p* < 10^-4^, Table 1).

**Table 1.** Comparison between ASE and Non-ASE genes

| Category | ASE (n=418) | Non ASE (n=465) | P-value |
| --- | --- | --- | --- |
| Gene length (Kb) | 58 ± 4 | 63 ± 3 | 0.31 |
| 5' UTR length (Kb) | 0.23 ± 0.03 | 0.25 ± 0.01 | 0.15 |
| 3' UTR length (Kb) | 1.80 ± 0.08 | 1.87 ± 0.06 | 0.54 |
| Transcripts (coding) per gene | 1.50 ± 0.05 | 1.65 ± 0.06 | 0.10 |
| Exons per transcript | 12.4 ± 0.37 | 12.9 ± 0.29 | 0.31 |
| Cis-regulatory elements per Kb | 0.37 ± 0.01 | 0.34 ± 0.01 | 0.11 |
| Transcription factor binding sites per Kb | 1.61 ± 0.07 | 1.70 ± 0.07 | 0.38 |
| Sequence variants per Kb | 8.20 ± 0.25 | 4.59 ± 0.02 | <0.001 |
| Paralogs per gene | 5.31 ± 0.36 | 3.56 ± 0.22 | <0.001 |
| Evolutionary conservation (GERP++ score) per Kb | 201.33 ± 10.85 | 274.15 ± 13.26 | <0.001 |

We also compared promoter complexity. There are no differences in the density of liver-specific cis-regulatory elements defined using mouse ENCODE data (STAMATOYANNOPOULOS *et* al. 2012). Similarly, there are no differences in the density of transcription factor binding sites (TFBS) defined using a comparative genomic approach (DAILY *et al.* 2011) (Table 1). However, the subset of genes with no or low ASE are enriched (*p* < 10^-46^, hypergeometric test) in housekeeping genes (EISENBERG and LEVANON 2013). In fact, nearly 50% of the non-ASE set are house-keeping genes. In contrast only 20% of the ASE set belong to this category.

Another distinguishing characteristic of the two sets is their density of sequence variants. The mean density in the non-ASE set is significantly different from the ASE set (4.59 ± 0.02 versus 8.20 ± 0.25 per Kb, p ∼ 0.0, two-tailed *t* test). This marked difference suggests that genes in the non-ASE set are under comparatively stronger purifying selection.

To test whether ASE and non-ASE sets are subject to different levels of purifying selection (the elimination of deleterious sequence variants) we compared the strength of selective constraint (GERP++ scores) on genomic regions across 33 mammalian species (COOPER *et al.* 2005; DAVYDOV *et al.* 2010). The ASE gene set (201.33 ± 10.85) have significantly lower conservation scores than the non-ASE set (274.15 ± 13.26) indicating that they tolerate and accumulate more mutations; a subset of which are highly likely to modulate expression (*p* < 10^-4^, two-tailed *t* test, Table 1).

To evaluate if genes in ASE and non-ASE sets belong to different functional categories, we compared them for overrepresented gene ontology and KEGG pathway terms using DAVID functional annotation tool (HUANG DA *et al.* 2009). ASE genes are significantly enriched (Benjamini corrected *p* < 0.05) in genes associated with KEGG pathway terms including ‘complement and coagulation cascades’, ‘retinol metabolism’, ‘metabolism of xenobiotics by cytochrome P450’, and ‘lipid, fatty acid and steroid metabolism’. Non-ASE genes are enriched in genes with gene ontology (GO) terms representing broad functional categories such as ‘macromolecule localization’, ‘catalytic activity’, and ‘ubiquitin mediated proteolysis’.

To evaluate the effect of ASE on phenotypes we performed phenotype enrichment analysis (WENG and LIAO 2010) on mouse-mutant phenotypes (BLAKE *et al.* 2009) derived from Mouse Genome Informatics (MGI, www.informatics.jax.org/phenotypes.shtml). As noted above, ASE genes compared to non-ASE genes are enriched (unadjusted *p* < 0.01) for phenotypes including ‘abnormal gall bladder physiology’ (MP:0005085), ‘abnormal xenobiotic induced morbidity/mortality’ (MP:0009765), and ‘abnormal glucose homeostasis’ (MP:0002078).

### Identification of *cis* eQTLs

We performed linkage-based eQTL mapping using a gene expression data set generated using liver samples from 40 BXD strains (Methods). Of the ∼45,000 probe sets, we selected ∼41,500 that have a uniquely assigned gene identifier. This subset represents ∼19,000 genes. *Cis* eQTLs were required to have LOD scores greater than 3 and LOD peaks within ± 5 Mb of their cognate gene. A LOD score ≥ 3 roughly corresponds to a nominal *p* value of < 0.001 and is widely used to indicate a high probability of linkage. We detected a total of 1,907 *cis* eQTLs corresponding to 1,474 genes (Fig. 2, Table S8). *cis* eQTLs with very high LOD scores (≥ 25) include *Snx6, Adi1, Cfh, Fbxo39,* and *St3gal4*.

### Variant overlapping probes cause spurious *cis* eQTLs

SNPs and indels in probe sequences can influence hybridization kinetics and cause incorrect measurement of expression. Twenty-five percent of apparent *cis* eQTLs detected in the hippocampus are probably caused by variants in probes rather than by genuine differences in expression (CIOBANU *et al.* 2010). We identified 739 cis-modulated probe sets that overlap *D* variants. To evaluate how these variants affect the direction and size of additive effects for corresponding *cis* eQTLs, we compared *cis* eQTLs between probe sets with variants and probe sets without variants. Probe sets without variants were precisely balanced with respect to *B* versus *D* effects. In contrast probes with variants were highly imbalanced and ∼70% were associated with high *B* expression. Of 408 *cis* eQTL genes represented by probes with variants, 193 could be compared with results from ASE analysis, and 149 genes showed the same direction of expression bias as ASE. A total of 1,215 genes were associated with *cis* eQTLs.

### Cis-modulated genes from ASE and eQTL mapping overlap

We compared results from ASE with those from eQTL mapping. Of the 3,431 genes that were jointly tested, 1,808 (∼50%) and 867 (∼25%) were identified as cis-modulated by ASE and eQTL mapping, respectively. Six hundred and eighty-three genes were jointly identified as cis-modulated, a significant overlap (hypergeometric *p* < 10^-73^), and ∼90% had the same effect polarity (Table S9). One thousand one hundred and twenty-five and 184 cis-modulated genes were exclusively identified by ASE and eQTL mapping respectively. In other words, roughly 80% of *cis* eQTLs also have ASE differences and ∼40% of ASE differences are associated with *cis* eQTLs. To investigate discrepancies, we compared LOD scores of jointly identified cis-modulated genes with those only identified by eQTL mapping. The joint set exhibit significantly higher LOD scores (*p* < 10^-5^) (Fig. 4). We further compared the RNA-seq read depth (expression) of these two groups and the joint set has significantly higher expression (three-fold difference, *p* = .03, Fig. 5). This suggests that the ASE analysis lacks adequate read depth (statistical power) to detect allelic differences corresponding to *cis* eQTLs with comparatively low LOD scores. We performed an empirical power analysis (Fig. 6) to illustrate dependency of ASE analysis on read-depth to detect allelic differences of different magnitudes. As expected, strong differences can be reliably detected with a relatively small number of reads and vice-versa. The joint set also has higher allelic expression differences compared to 1,125 genes identified only by ASE (*p* < 10^-30^) (Fig. 7). We used a stringent LOD threshold ≥ 3 to define *cis* eQTLs and this will reduce the number of *cis* eQTLs corresponding to genes with low ASE. We therefore performed single marker analysis at less stringent FDR < 0.1 (see Methods) to identify cis-modulated genes and compared them with the ASE gene set. The number of jointly identified genes increased from 648 (eQTL mapping, LOD ≥ 3) to 962 (single marker analysis, FDR < 0.1), and those exclusively identified by ASE were reduced from 1,125 to 774. Thus a large fraction of disjoint between ASE and eQTL results are explained by the different statistical criteria we used to define both ASE genes and *cis* eQTLs.

**Figure 4.**
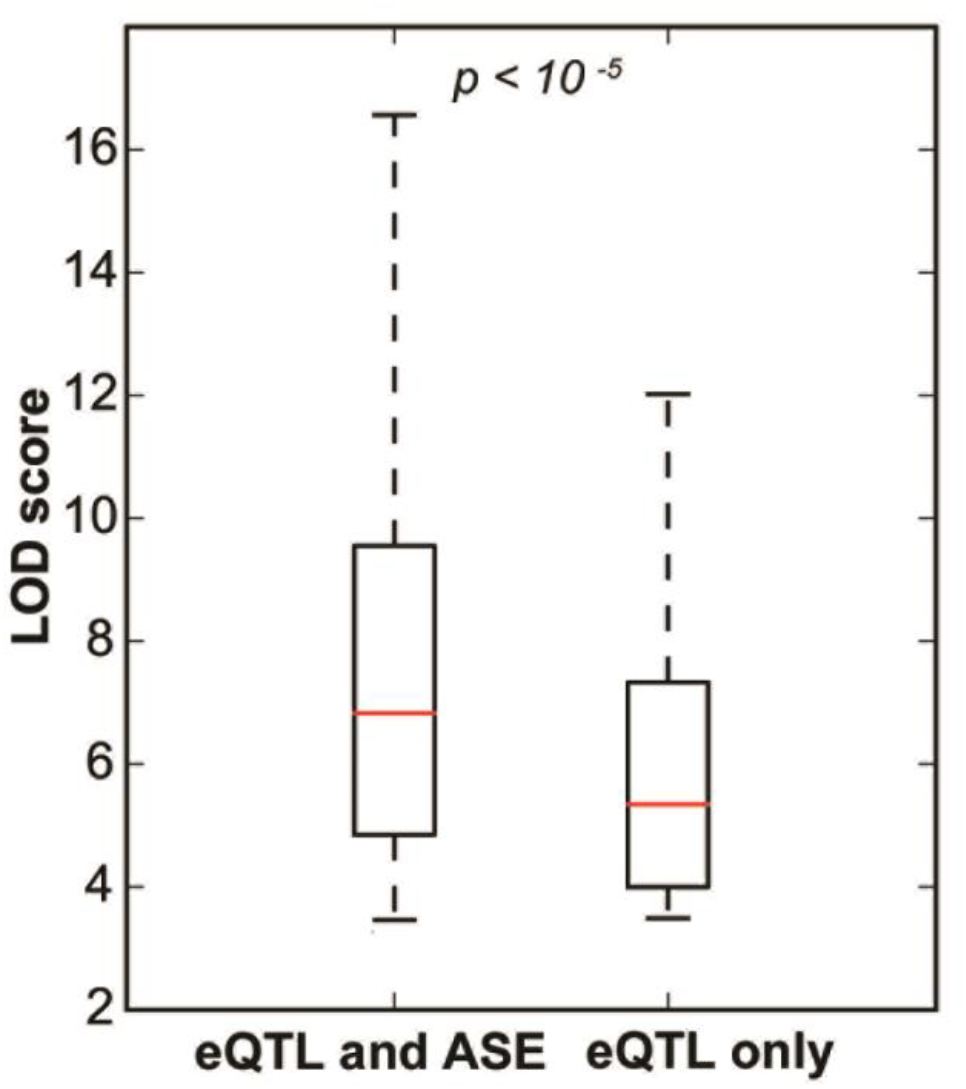
Comparison of LOD scores from jointly identified cis-modulated genes (ASE and eQTL mapping, left boxplot) with those only identified using eQTL mapping (right boxplot). The *Y*-axis represents LOD scores. Outliers are not shown.

**Figure 5.**
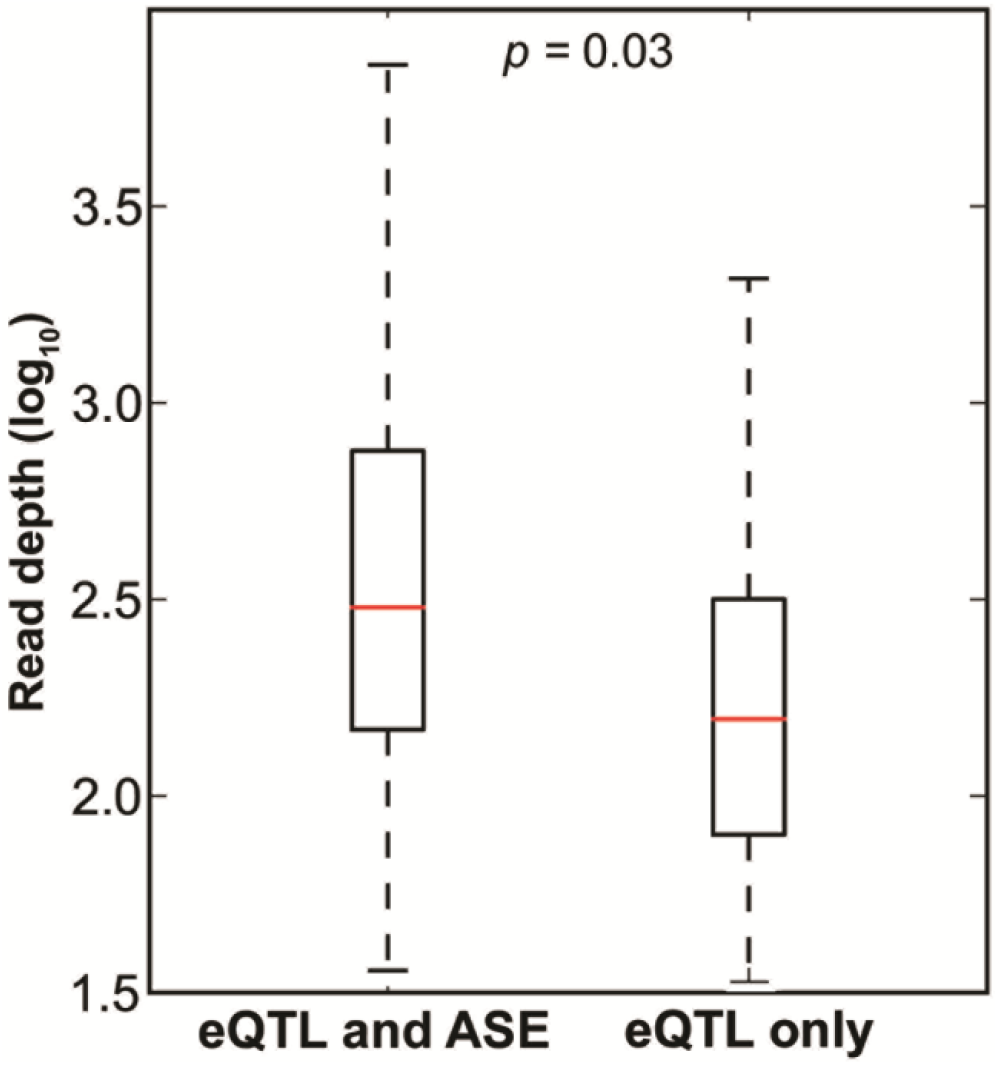
Comparison of RNA-seq read depth (log_10_) from jointly identified cis-modulated genes (ASE and eQTL mapping) with those only identified using eQTL.

**Figure 6.**
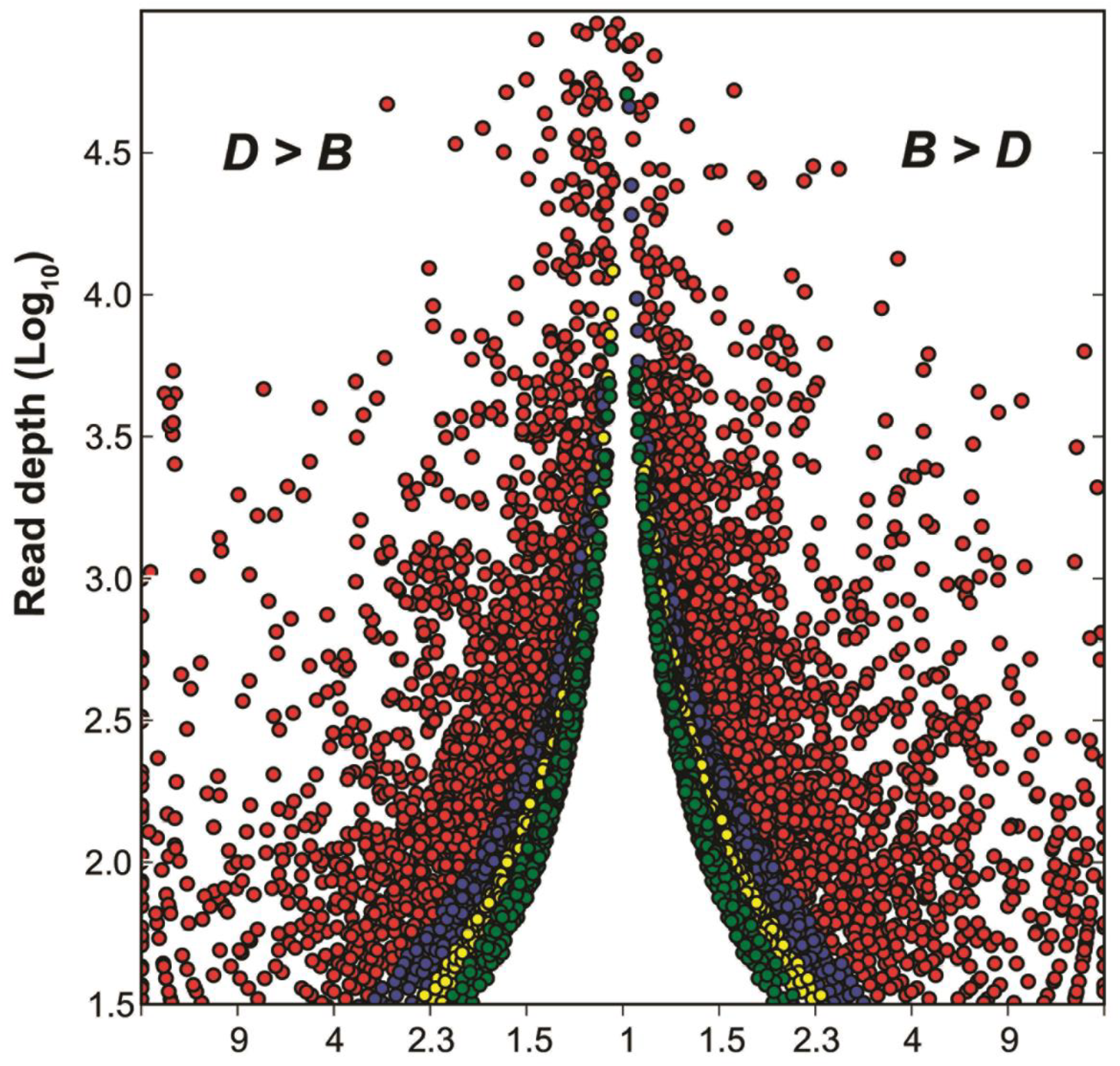
Empirical determination of read depth required to detect allelic differences of a given size. Each circle represents a SNP. The *X*-axis represents the measured fold-difference and the *Y*-axis represents RNA-seq read depth (log_10_) for a given SNP. SNPs exhibiting ASE at an FDR threshold of ≤ 0.2 have been plotted as circles. Red circles represent SNPs with ASE at an FDR threshold of ≤ 0.01. The red and blue circles, combined, represent SNPs with ASE at an FDR threshold of ≤ 0.05. Similarly, red, blue and yellow circles, combined, represent SNPs with ASE at an FDR threshold of ≤ 0.1.

**Figure 7.**
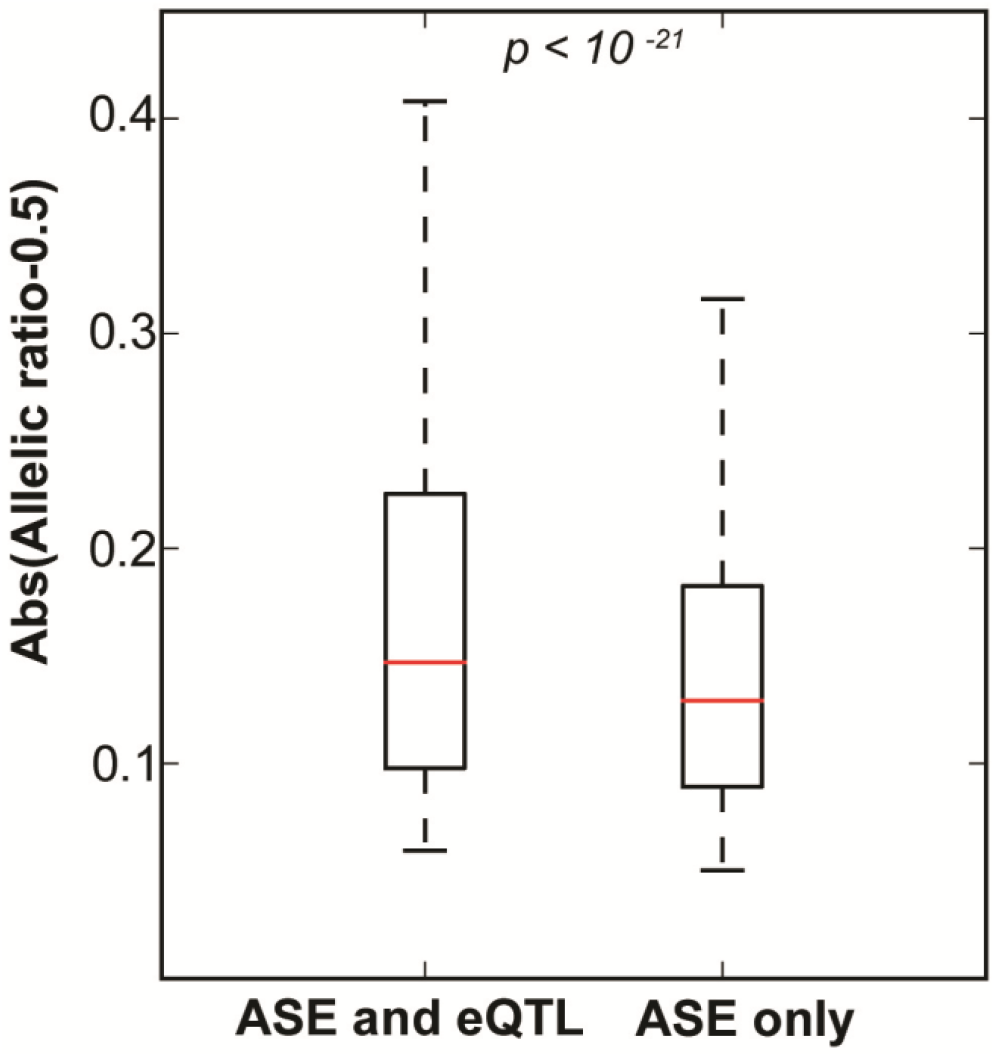
Comparison of absolute allelic differences from jointly identified cis-modulated genes (ASE and eQTL mapping, left boxplot) with those only identified using ASE (right boxplot). Outliers are not shown.

### Genetic variants affecting transcript abundance and protein abundance show poor overlap

We performed linkage based protein QTL (pQTL) mapping on liver proteomics data generated from a set of 38 BXD strains (WU *et al.* 2014). One hundred and seventy-two autosomal proteins involved in metabolism were quantified using a targeted mass spectrometry method. Only 7% (*n* = 12) are associated with *cis* pQTLs, including ABCB8, ACADS, ACOX1, ATP5O, BCKDHB, CAR3, DHTKD1, GCLM, MRI1, NNT, PM20D1, and TYMP (Fig. 2, where a *cis* pQTL must have a LOD > 2 located within ±5 Mb of the parent gene). Not surprisingly, all of the *cis* pQTLs are also associated with significant ASE differences with matched polarity (Table S10).

Similarly, 8 of these *cis* pQTLs are linked to *cis* eQTLs with high LOD scores and with matched polarity. However, 39 genes with significant ASE and 18 genes (Fig. 8) with significant *cis* eQTL are not associated with *cis* pQTLs. For example, *Ddah1* has significant ASE (3–4 fold difference) and a strong *cis* eQTL (chr3:145 Mb, LOD ∼ 14.5) favoring the *B* allele. However, the protein difference across BXDs does not map to the location of gene and protein difference between *B* and *D* alleles has a one-tailed *p* of 0.2—a reasonably strong negative result. This case is doubly interesting because variation in DDAH1 protein maps as a *trans* pQTL (Chr7: 27.85 Mb, LOD ∼ 2.5).

**Figure 8.**
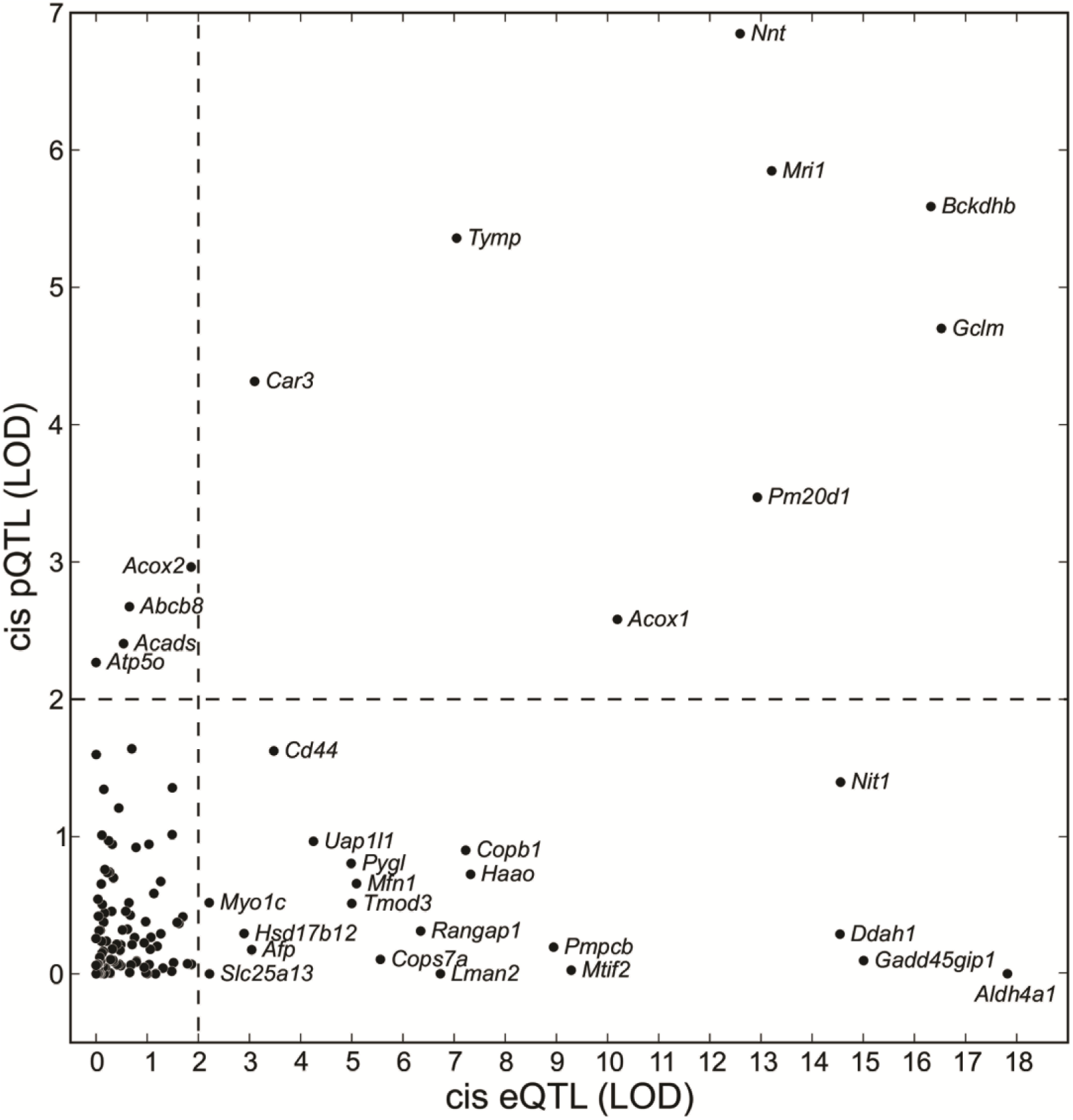
Comparison of cis-acting variation at transcript versus protein levels. The *X*-axis and *Y*-axis represent LOD scores for genes and cognate proteins measured at their closest markers (*cis* LOD). A LOD of 2 (dashed line) roughly corresponds to a nominal *p* < 0.01.

### Majority of aberrant alleles do not affect expression severely

Nonsense mediated decay (NMD) is a molecular surveillance mechanism that selectively degrades aberrant transcripts produced as a result of nonsense or splice-site variants (BAKER and PARKER 2004; LAREAU *et al.* 2007; ZHANG *et al.* 2009). NMD of aberrant transcripts should result in extreme allelic ratios (close to zero or one). However, over two-thirds of nonsense variants (transcripts) in human cell lines escape NMD through unknown mechanisms (LAPPALAINEN *et al.* 2013). We measured allelic ratios for 12 nonsense variants (transcripts) and remarkably only two—*Gbp11* (0.05, high *D*) and *Mug2* (0.90, high *B*)—had extreme ratios across multiple SNPs. Interestingly, half of the stop codon losses identified in the *B* allele only add one to two amino acids to the variant proteins, including VMN2R79 (+1 amino acid), ADAM3 (+1), SPNS3 (+2), ZCCHC9 (+2), DLGAP5 (+2), and HOGA1 (+1). None of these transcripts have extreme allelic ratios. We found that a third of murine genes have one or more in-frame stop codons in close vicinity (<30 nucleotides) to the original stop codon. Tandem stop codons are also known to be conserved in yeast (LIANG *et al.* 2005), and may provide a safeguard against stop codon losses. We also evaluated 36 splice-site variants and only five of these transcripts, including *Cyp2c39* (0.02), *Arhgef10* (0.03), *Pik3c2g* (0.05), *Lox14* (∼0.9), and *Rpsa* (0.99) had extreme ratios. Splicing machinery may also use alternative splice sites in the close vicinity to the original splice site to prevent the production of aberrant transcripts. For example, we found a polymorphism (rs33609674) within intron 1 of *Hcfc1r1* that introduces a CAG splice acceptor site in the *B* allele. The acceptor site is located twelve nucleotides upstream of exon 2 of the *D* allele (Fig. S2) resulting in the addition of four amino acids and presumably no seriously aberrant transcript is produced in either case.

In conclusion a majority (> 85%) of presumed aberrant transcripts including nonsense and splice-site variants escape NMD. We speculate that the use of alternative stop codons or splice sites in the immediate vicinity of the primary mutation apparently prevents aberrant transcript production.

### Mechanistic insights into the basis of allele-specific expression—quantitative and qualitative differences

*Cis*-acting variants affect expression in three major ways: (1) by modulating transcription rates and stability (mRNA abundance), (2) by modulating transcript processing (splicing and polyadenylation), and (3) by altering mRNA transport and storage (AN *et al.* 2008). Allelic ratios of SNPs that represent different regions of a transcript can be collectively analyzed and compared to provide mechanistic understanding of these alternative mechanisms. Multiple SNPs that have the same polarity and roughly the same magnitude of effect suggest variants in enhancers or transcription-factor binding sites that control transcript levels globally. For example, *Nnt*, a gene linked to insulin hypersecretion in the *D* parent (ASTON-MOURNEY *et al.* 2007), has a strong *cis* pQTL (LOD ∼8) in liver with high expression of the *D* allele. All eight SNPs exhibit significant ASE and with the same polarity (Fig. 9A). Another example is *Gclm*, a gene involved in the metabolism of dietary lipid (KENDIG *et al.* 2011), that also has a strong *cis* pQTL (LOD ∼5) in liver with high expression of the *B* allele. All five SNPs have ASE with the same polarity.

**Figure 9.**
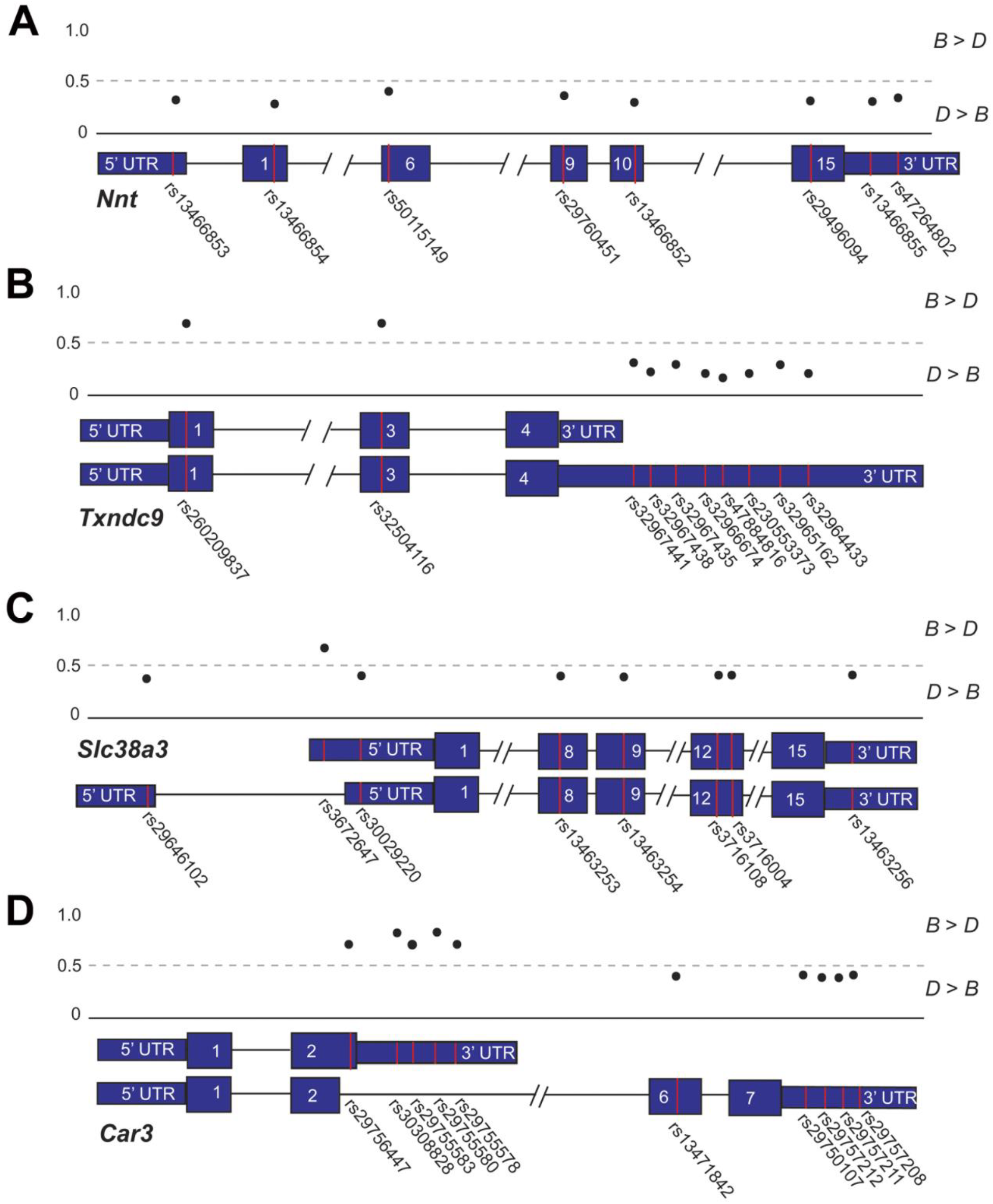
Schematic examples of genes potentially associated with different categories of *cis*-regulatory mechanisms. The *Y*-axis shows the allelic ratios of SNPs located within the gene. An allelic ratio greater than 0.5 (dashed line) represents high expression of the *B* allele. Examples of allele-specific regulation of (A) overall gene expression, (B) 3’ UTR processing, (C) 5’ UTR processing, and (D) isoform usage.

Neighboring SNPs located within 5’ or 3’ UTRs that have opposite polarity suggest allele-specific differential usage of alternate transcriptional initiation or polyadenylation sites. One example is *Txndc9*, a gene linked with colorectal cancer in humans (LU *et al.* 2012). This gene has multiple transcripts with alternative polyadenylation sites as demonstrated by multiple mRNAs in RefSeq and Ensembl gene models. Two SNPs located in exons 1 and 3 have significantly ASE with high *B* expression whereas eight SNPs located in the extended 3’ UTR (Fig. 9B) have high *D* expression suggesting allele-specific differences in 3’ UTR processing. A similar pattern is observed in array data: probe sets in coding exons have high *B* expression whereas those in the 3’ UTR have high *D* expression. The longer 3’ UTR of the *D* allele harbors putative binding sites (PhastCons > 0.5 and mirSVR score < -0.3 (BETEL *et al.* 2010)) for multiple miRNAs, including *miR-539*, *miR-96*, *miR-129-5p*, that may explain overall low expression of the *D* transcript. Another example is *Slc38a3*, a glutamine transporter involved in ammonigenesis (BUSQUE *et al.* 2014; BUSQUE and WAGNER 2009). GenBank and Ensembl gene models demonstrate multiple transcripts that use alternative transcriptional initiation sites. Four SNPs in exons and a SNP in 3’ UTR have high *D* expression. However, SNPs in the 5’ UTR have variable ASE (Fig. 9C). Two SNPs (rs30029220 and rs29646102) exclusive to the longer 5’ UTR have high *D* expression whereas a SNP (rs3672647) in the shorter 5’ UTR has high *B* expression suggesting that the *D* allele favors usage of transcript with the longer 5’ UTR.

Finally, SNPs with opposite ASE polarity in different coding exons are probably caused by alternative exon usage or alternative splicing. For example, carbonic anhydrase 3 (*Car3*), a gene linked to adipogenesis (MITTERBERGER *et al.* 2012), has a strong *cis* pQTL (LOD ∼5) with high expression of the *D* allele. Five SNPs located exclusively in a long isoform show significant ASE with high *D* expression: one SNP in the exon 6 and four SNPs in the 3’ UTR (Fig. 9D). In contrast, 5 SNPs located exclusively in the short isoform have high *B* expression.

## DISCUSSION

Allele-specific expression differences are a major driver of phenotypic differences and variation in disease risk. We exploited RNA-seq and eQTL data sets to quantify the extent and intensity of cis-acting variation in expression in liver. After correcting for alignment bias, we achieved the expected symmetrical distribution of allelic differences. Well separated SNPs within single exons are highly concordant both in strength and polarity of effects. Allelic ratios of SNPs also correlate well across biological replicates. The concordance between ASE and *cis* eQTLs is strong.

Having dealt with these technical challenges, we were able to identify statistically significant ASE differences with minimum fold difference of 1.25x for nearly half of all assayed transcripts. This latter finding strongly supports recent work by Crowley and colleagues (CROWLEY *et al.* 2015) demonstrating pervasive and high levels of ASE in brain and other tissues. In each F1 strain contrast, they detected significant ASE in 50% or more of all tested genes/transcripts at an FDR of 0.05. In total, 90% of testable genes exhibited ASE effects in at least one pair of strains. Lagarrigue and colleagues (LAGARRIGUE *et al.* 2013) detected somewhat less pervasive ASE effects (∼20%) in liver of C57BL/6JxDBA/2J F1 animals, but this is most certainly a matter of lower RNA-seq read depth (statistical power), and a higher fold difference (1.5x) criterion they used to identify ASE. Of 2,256 genes, they only observed 383 genes with significant ASE. We are now able to address three questions posed in the introduction.

### Highly conserved genes have low levels of ASE

Do differences in the magnitude of ASE represent differences in complexity of expression control or in evolutionary history? To answer these questions we compared a group of genes with very low and very high ASE. We found no differences in the density of cis-regulatory elements, but genes with low ASE do appear to be under more intense purifying selection. Fifty percent of the non-ASE set are house-keeping genes (EISENBERG and LEVANON 2013) and are likely to evolve comparatively slowly (She et al. 2009; Zhang and Li 2004). In contrast, genes with high ASE are likely to have higher functional redundancy as estimated indirectly by numbers of paralogs, and they are also enriched in tissue-specific functions. In our study of liver they are involved in the metabolism of lipids, fatty acids, and xenobiotics. We speculate that high gene sets with higher ASE may function in tissue-specific pathways that tend to retain both higher numbers of paralogs and be under less evolutionary constraint. The comparatively high range of variation in expression of these genes may be crucial to conferring greater physiological tolerance to noise and environmental challenges. ASE may also be one of the genetic mechanisms that underlie the canalization of phenotypes (MASEL and SIEGAL 2009; WADDINGTON 1942).

### Genetic variants affecting transcript abundance and protein abundance show poor overlap

Transcript abundance has been shown to correlate only modestly with protein abundance. (ALBERT *et al.* 2014; GHAZALPOUR *et al.* 2011; MAIER *et al.* 2009; SKELLY *et al.* 2013; WU *et al.* 2014) and we add the corollary that genetic variants that affect transcript abundance and protein abundance show low concordance. As expected, there is considerable disparity in allelic variation detected at mRNA and protein levels (Fig. 8). A few of the *cis* eQTLs with very high LOD scores (≥10) but essentially no *cis* pQTLs (LOD score < 1) are *Ddah1*, *Gadd45gip1* and *Aldh4a1*. Fu and colleagues suggested an increased buffering at the level of proteins and metabolites, such that only a few genetic variants modulate major phenotypic variation and majority of them remain silent (FU *et al.* 2009). Factors that are known to contribute towards the disparity between mRNA and protein levels include post-transcriptional and post-translational modifications (MAIER *et al.* 2009), differences in half-lives (SCHWANHAUSSER *et al.* 2011), variability in mRNA expression level due to changes in cell-cycle (CHO *et al.* 1998).

### Comparison between cis-modulated genes identified by ASE and eQTL mapping

We found significant overlap in cis-modulated genes identified by ASE and eQTL analysis, despite substantial differences between methods and assays. Eighty percent of *cis* eQTL genes are also detected by ASE, and ∼90% of them have the same polarity. The set of 683 genes identified by both methods have significantly higher LOD scores (Fig. 4) than the set of 184 genes identified only by eQTL mapping. In our work, when ASE methods fail to detect known eQTLs, this is almost certainly due to inadequate read depth (statistical power). High sampling error in RNA-seq data will affect power of ASE analysis especially for genes with low expression (PANDEY and WILLIAMS 2014; TARAZONA *et al.* 2011). As shown in Fig.6, high read depth is required to detect small allelic difference. A small fraction of presumed *cis* eQTLs can be local trans eQTL effects of neighboring genes.

The jointly identified set of ∼650 genes has greater allelic differences than the set of 1,125 genes identified only by ASE (Fig. 7). A large fraction of subtle allelic differences identified only by ASE may have been confounded by noise or epistatic trans-acting effects in the eQTL analysis. The small sample size of the BXD cohort used for the eQTL mapping may not have adequate statistical power to map weak *cis* eQTLs, especially in the presence of epistatic *trans* eQTLs. Additionally, the LOD threshold of greater than 3 used to define cis eQTLs may be too stringent in this particular context.

To the best of our knowledge, Babak and colleagues (BABAK *et al.* 2010) were the first to compare F1-derived ASE results with eQTL results from F2 intercrosses for adipose and islets samples. They found an 80% overlap between genes exhibiting ASE and genes with *cis* eQTLs. Lagarrigue and colleagues (LAGARRIGUE *et al.* 2013) found a 60% overlap between the methods. Hasin-Brumshtein and colleagues (HASIN-BRUMSHTEIN *et al.* 2014) performed a similar comparison in adipose tissue and reported relatively poor overlap (∼20%), but as noted above, differences with our more concordant results are most likely cumulative result of differences in criteria and ratios of statistical power of ASE analysis using F1 hybrids and cis eQTLs analysis using large intercrosses.

As highlighted in the introduction, it is now clear that most common variation in phenotype and disease risk are linked to variants that modulate patterns of gene expression. ASE is a sensitive and a cost-effective method to detect cis-acting differences in expression. Environmental and trans-acting factors are fully controlled in isogenic F1 individuals, and ASE analysis only requires a small F1 sample size. In this respect it has a clear advantage over eQTL analysis of segregating populations. However, many classical laboratory strains have been derived from ancestral stock with limited haplotype diversity. As a result, a large fraction of an F1 genome will be identical by descent (IBD) and genes in these regions cannot be interrogated using ASE.

Linkage-based eQTL analysis adds two important dimensions to an ASE study. First, it makes it possible to assign causality to specific variants using high-resolution mapping populations (LI *et al.* 2010; WANG *et al.* 2012). Second, eQTL analysis makes it possible to study the downstream effects of differential expression. These downstream effects are detected as *trans* eQTLs of other mRNAs or proteins.

### ASE varies between different environments and genetic backgrounds

Estimates of ASE and *cis* eQTL will vary as a function of genetic background (CROWLEY *et al.* 2015), tissue (KEANE *et al.* 2011), environment (WU *et al.* 2014), and sex (MOZHUI *et al.* 2012). For example, cis eQTLs effects can be strongly dependent on diet. The *cis* eQTL associated with *Ndusf2* increases from LOD score of 2 in a mouse cohort on a normal chow diet to a LOD score of 6 in a cohort on a high fat diet (Wu et al. 2014). For these reasons, one should not expect estimates of ASE in liver of one population or treatment to generalize. Nevertheless, many of the large ASE effects caused by strong cis-acting variants will often be well conserved across environments, cell types, and genetic backgrounds. For example, ASE effects due to copy number variants (DISTLER and PALMER 2012), retrotransposons disrupting 3’ UTRs (LI *et al.* 2010), and nonsense mutations (WILLIAMS *et al.* 2014) will often produce strong and consistent ASE effects across many tissues and treatments.

## MATERIALS AND METHODS

### Genomic data for DBA/2J

We downloaded paired-end sequencing data for DBA/2J from the European Nucleotide Archive, accession number ERP000044 (KEANE *et al.* 2011). Data consists of nine paired-end libraries sequenced on the Illumina GAII. Read lengths vary between 54–76 nt. We also downloaded Illumina paired-end sequencing data from the Sequence Read Archive, accession number SRP001135 (WANG *et al.* 2010). These data consists of three libraries sequenced on the GAII with read lengths of 100 nt.

### DBA/2J genomic read alignment and variant calling

Reads were trimmed to remove low quality bases and aligned to the C57BL/6J reference genome (mm10) using Burrows Wheeler Aligner (BWA) (LI and DURBIN 2010) (version 6.1). Base quality scores were recalibrated at the lane level using Genome Analysis Toolkit (GATK, v2.7) ‘TableRecalibration’ (MCKENNA *et al.* 2010). All lanes for each library were merged into one BAM file using Picard (version 1.8, http://picard.sourceforge.net/) and duplicates were flagged using ‘MarkDuplicates’. BAM files for each library were combined together to create a single master BAM file containing all *D* sequences. Finally, GATK ‘IndelRealigner’ was used to realign reads near indels from the Mouse Genome Project (KEANE *et al.* 2011), as well as potential indels predicted by GATK. SNPs and indels were identified using the GATK Unified Genotyper and the default settings. An in-house Python script was used to remove low quality variants based on multiple criteria including strand bias, minimum read mapping quality, end distance bias, minimum and maximum depth, and proximity to indels (https://github.com/ashutoshkpandey/Variants_call/blob/master/Filter_GATK_vcf.py). Only homozygous SNPs and indels were retained. For CNV detection, we used copy number detector (cnD) at the default settings (SIMPSON *et al.* 2010). We used SnpEff (CINGOLANI *et al.* 2012) to annotate SNPs and indels against RefSeq gene models, and categorized them as nonsense, splice site, frameshift, or missense.

### RNAseq data for C57BL/6JxDBA/2J hybrids

We downloaded paired-end RNA-seq data from the European Nucleotide Archive (accession number ERP000591) for liver of C57BL/6JxDBA/2J F1 female hybrids generated by crossing C57BL/6J females with DBA/2J males (KEANE *et al.* 2011). The data consist of transcriptome sequence from six biological replicates. We acquired a total of ∼181 m read pairs (2x76 nt in length). We removed low quality reads and used the remaining ∼173 m read pairs for alignment.

### RNA-seq read alignment

We aligned RNA-seq reads to both the C57BL/6J reference genome (mm10 assembly) and the DBA/2J genome using “Splice Transcripts Alignment to a Reference” tool (STAR, version 2.3.1a) (DOBIN *et al.* 2013) with the following parameters “--outFilterMultimapNmax 10 -- outFilterMismatchNmax 12”. Read pairs that were not aligned in concordance with the library design, in particular read strand, were removed. We allowed a maximum insert size of 300,000 nucleotides (maximum intron length) to allow alignment of those read-pairs aligned to different exons. We selected read pairs for which both reads were uniquely aligned and for which each had less than six mismatches. If one member of a read-pair could not be aligned then we retained the other member only if it could be aligned uniquely.

### Calculation of allelic ratio

We used SAMtools (version 0.1.19) “pileup” function (LI *et al.* 2009) and an in-house Python (https://github.com/ashutoshkpandey/ASE_prealignment/blob/master/Allele_specific_SAM.py) script to assign reads to their parental allelic origin by comparing alignments to the C57BL/6J and the SNP-substituted DBA/2J genome. If reads were aligned to both genomes then we required them to map at the same locations. Those reads that overlapped SNPs were assigned to their parental allele origin. To ensure that differential expression was not due to amplification by PCR during library preparation, we removed all potential PCR duplicates except for the single read with the fewest mismatches using Picard’s MarkDuplicates tool (version 1.78). We calculated allelic ratios for each SNP defined as the ratio of number of reads assigned to the reference allele (*B*) to the total number of aligned reads (*B*+*D*).

### Definition of ASE using chi-square test

For each SNP we used an interquartile range (IQR) method to identify outlier allelic ratios from the set of F1 replicates. Outlier ratios were located outside the [Q1 – 1.5(IQR) and Q3 + 1.5(IQR)] range where Q1 and Q3 represent first and third quartiles and IQR is calculated as Q3 – Q1. Reads from replicates showing concordant allelic ratios were merged and allelic ratios were recalculated. We used the chi-square goodness of fit test to determine allelic imbalances for a given SNP. For a SNP showing an allelic imbalance, the ratio will deviate from 0.5. We defined genes as having an allele-specific expression difference if they contained one or more SNPs with an allelic imbalance at an FDR threshold of less than 0.1 (BENJAMINI and HOCHBERG 1995). We also required the expression fold difference to be >1.25.

### Array expression data and eQTL mapping

We used an Affymetrix data set (Mouse 430 v2.0 array) consisting of liver gene expression data for 40 genetically diverse BXD strains (GeneNetwork.org accession GN310, http://genenetwork.org/webqtl/main.py?FormID=sharinginfo&GN_AccessionId=310). We performed robust multichip analysis (RMA) preprocessing and rescaled values to log_2_ and stabilized the variance across samples (GEISERT *et al.* 2009). We used QTL Reaper, mapping code that uses the method of Haley and Knott for eQTL analysis (HALEY *et al.* 1994), and a set of 3,200 markers. We excluded probe sets located on X and Y chromosomes (∼2,500 probe sets). Locations of probe sets were identified using custom annotation files. Similarly, we performed pQTL mapping on expression data from 172 proteins (WU *et al.* 2014). This data can be downloaded from Genenetwork.org (accession GN490, http://www.genenetwork.org/webqtl/main.py?FormID=sharinginfo&GN_AccessionId=490). To identify Affymetrix probes that overlapped sequence variants, we first aligned probe sequences against the mouse reference genome (mm10) using BLAT (KENT 2002), and then compared genomic coordinates of probes for overlap with sequence variants.

### Comparison between ASE and non-ASE genes (URLs)

We downloaded the liver-specific regulatory elements data from Ensembl Regulatory build (ftp://ftp.ensembl.org/pub/release-81/regulation/mus_musculus); see more details on this build here: http://www.ensembl.org/info/genome/funcgen/regulation_sources.html. For TFBS comparison we used data from MotifMap—genome-wide maps of regulatory elements. The file was downloaded using the following link: (http://www.igb.uci.edu/∼motifmap/motifmap/MOUSE/mm9/multiz30way/MotifMap_MOUSE_mm9.multiz30way.tsv.bz2). A list of house-keeping genes was downloaded using the following link: http://www.tau.ac.il/∼elieis/HKG. In order to compare for the evolutionary conservation between the ASE and non-ASE genes, we used GERP++ scores for mouse (http://mendel.stanford.edu/SidowLab/downloads/gerp/mm9.GERP_elements.tar.gz). We downloaded *M. musculus* and *H. sapiens* paralog data from Ensembl BioMart (www.ensembl.org/info/data/biomart.html) (HAIDER *et al.* 2009; VILELLA *et al.* 2009). All the mm9 coordinates were converted to mm10 using UCSC liftOver utility. The counts/scores of cis-regulatory elements, TFBSs, DBA/2J sequence variants, and GERP++ scores were normalized by gene length before comparison.

### Single marker analysis

We performed single marker analysis as an alternative to eQTL mapping to identify cis-modulation in expression. For each gene we selected its closest marker and classified BXDs by genotype (*B* allele or *D* allele) for that marker. We compared expression using a *t*-test and selected genes showing significant expression difference at an FDR of < 0.1.

## ACKNOWLEDGEMENT

We thank Drs. Robert Rooney and Divyen Patel (Genome Explorations Inc.) and Dr. Kristen Hamre (UTHSC) for allowing us to use their liver array data. We also thank Dr. Lu Lu and Dr. Megan Mulligan for their intellectual contribution to the study. This work supported by NIAAA Integrative Neuroscience Initiative on Alcoholism (U01 AA016662, U01 AA013499) and NIA (R01AG043930).

## FIGURES

**Figure S1.**
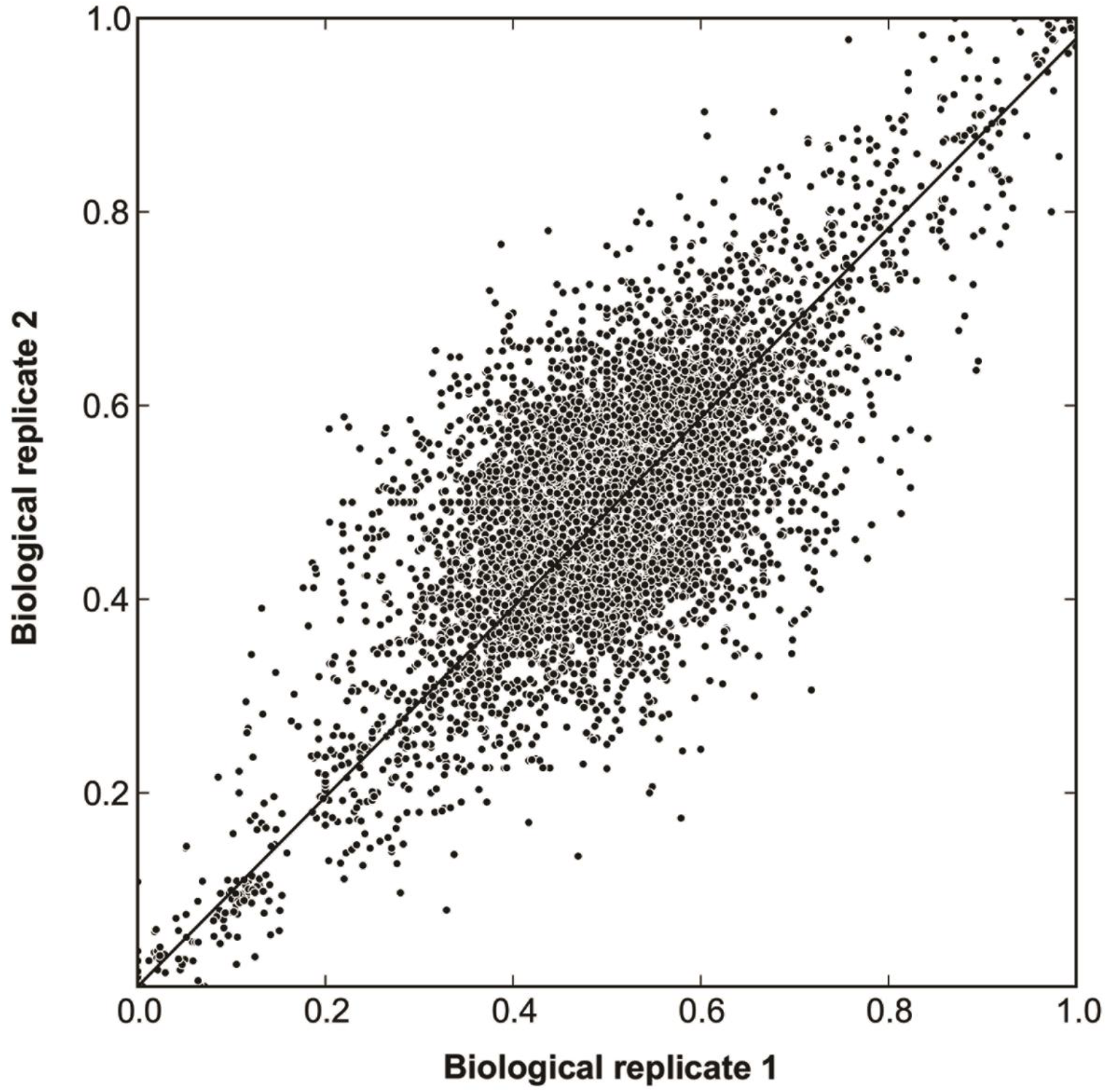
Pearson correlation of allelic ratios between two biological replicates (ERR032205 and ERR032206).

**Figure S2.**
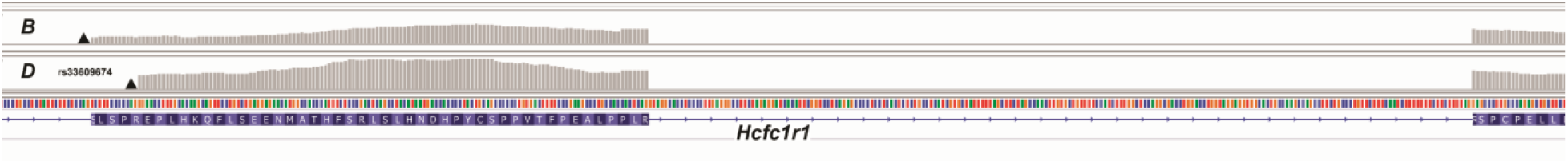
Alternative splice site usage between *B* and *D* alleles due to a polymorphism (rs33609674, C**A**G/C**G**G) that results in gain of splice acceptor site (solid triangle) in the *B* allele. The *B* allele uses an acceptor site (solid triangle) located in intron 1 of the *D* allele. As a result, the *B* transcript has 4 additional amino acids (SLSP) in the beginning of exon 2. Expression data from the striatum of the *B* (top track) and *D* (bottom track) alleles confirms the use of an alternate splice-site.

## REFERENCES

Albert, F. W., S. Treusch, A. H. Shockley, J. S. Bloom and L. Kruglyak, 2014 Genetics of single-cell protein abundance variation in large yeast populations. Nature 506: 494–497.

An, J. J., K. Gharami, G. Y. Liao, N. H. Woo, A. G. Lau et al., 2008 Distinct role of long 3' UTR BDNF mRNA in spine morphology and synaptic plasticity in hippocampal neurons. Cell 134: 175–187.

Andreux, P. A., E. G. Williams, H. Koutnikova, R. H. Houtkooper, M. F. Champy et al., 2012 Systems genetics of metabolism: the use of the BXD murine reference panel for multiscalar integration of traits. Cell 150: 1287–1299.

Aston-Mourney, K., N. Wong, M. Kebede, S. Zraika, L. Balmer et al., 2007 Increased nicotinamide nucleotide transhydrogenase levels predispose to insulin hypersecretion in a mouse strain susceptible to diabetes. Diabetologia 50: 2476–2485.

Babak, T., P. Garrett-Engele, C. D. Armour, C. K. Raymond, M. P. Keller et al., 2010 Genetic validation of whole-transcriptome sequencing for mapping expression affected by cis-regulatory variation. BMC Genomics 11: 473.

Baker, K. E., and R. Parker, 2004 Nonsense-mediated mRNA decay: terminating erroneous gene expression. Curr Opin Cell Biol 16: 293–299.

Bell, G. D., N. C. Kane, L. H. Rieseberg and K. L. Adams, 2013 RNA-seq analysis of allele-specific expression, hybrid effects, and regulatory divergence in hybrids compared with their parents from natural populations. Genome Biol Evol 5: 1309–1323.

Benjamini, Y., and Y. Hochberg, 1995 Controlling the false discovery rate: a practical and powerful approach to multiple testing. J R Stat Soc Series B Stat Methodol 57: 289–300.

Betel, D., A. Koppal, P. Agius, C. Sander and C. Leslie, 2010 Comprehensive modeling of microRNA targets predicts functional non-conserved and non-canonical sites. Genome Biol 11: R90.

Blake, J. A., C. J. Bult, J. T. Eppig, J. A. Kadin and J. E. Richardson, 2009 The Mouse Genome Database genotypes::phenotypes. Nucleic Acids Res 37: D712–719.

Brem, R. B., G. Yvert, R. Clinton and L. Kruglyak, 2002 Genetic dissection of transcriptional regulation in budding yeast. Science 296: 752–755.

Busque, S. M., G. Stange and C. A. Wagner, 2014 Dysregulation of the glutamine transporter Slc38a3 (SNAT3) and ammoniagenic enzymes in obese, glucose-intolerant mice. Cell Physiol Biochem 34: 575–589.

Busque, S. M., and C. A. Wagner, 2009 Potassium restriction, high protein intake, and metabolic acidosis increase expression of the glutamine transporter SNAT3 (Slc38a3) in mouse kidney. Am J Physiol Renal Physiol 297: F440–450.

Carneiro, A. M., D. C. Airey, B. Thompson, C. B. Zhu, L. Lu et al., 2009 Functional coding variation in recombinant inbred mouse lines reveals multiple serotonin transporter-associated phenotypes. Proc Natl Acad Sci U S A 106: 2047–2052.

Chesler, E. J., L. Lu, S. Shou, Y. Qu, J. Gu et al., 2005 Complex trait analysis of gene expression uncovers polygenic and pleiotropic networks that modulate nervous system function. Nature Genet 37: 233–242.

Cho, R. J., M. J. Campbell, E. A. Winzeler, L. Steinmetz, A. Conway et al., 1998 A genome-wide transcriptional analysis of the mitotic cell cycle. Mol Cell 2: 65–73.

Cingolani, P., A. Platts, L. WANG Le, M. Coon, T. Nguyen et al., 2012 A program for annotating and predicting the effects of single nucleotide polymorphisms, SnpEff: SNPs in the genome of Drosophila melanogaster strain w1118; iso-2; iso-3. Fly (Austin) 6: 80–92.

Ciobanu, D. C., L. Lu, K. Mozhui, X. Wang, M. Jagalur et al., 2010 Detection, validation, and downstream analysis of allelic variation in gene expression. Genetics 184: 119–128.

Cooper, G. M., E. A. Stone, G. Asimenos, E. D. Green, S. Batzoglou et al., 2005 Distribution and intensity of constraint in mammalian genomic sequence. Genome Res 15: 901–913.

Crowley, J. J., V. Zhabotynsky, W. Sun, S. Huang, I. K. Pakatci et al., 2015 Analyses of allele-specific gene expression in highly divergent mouse crosses identifies pervasive allelic imbalance. Nat Genet 47: 353–360.

Daily, K., V. R. Patel, P. Rigor, X. Xie and P. Baldi, 2011 MotifMap: integrative genome-wide maps of regulatory motif sites for model species. BMC Bioinformatics 12: 495.

Damerval, C., A. Maurice, J. M. Josse and D. De Vienne, 1994 Quantitative trait loci underlying gene product variation: a novel perspective for analyzing regulation of genome expression. Genetics 137: 289–301.

Davydov, E. V., D. L. Goode, M. Sirota, G. M. Cooper, A. Sidow et al., 2010 Identifying a high fraction of the human genome to be under selective constraint using GERP++. PLoS Comput Biol 6: e1001025.

Degner, J. F., J. C. Marioni, A. A. Pai, J. K. Pickrell, E. Nkadori et al., 2009 Effect of read-mapping biases on detecting allele-specific expression from RNA-sequencing data. Bioinformatics 25: 3207–3212.

Deveale, B., D. Van Der Kooy and T. Babak, 2012 Critical evaluation of imprinted gene expression by RNA-Seq: a new perspective. PLoS Genet 8: e1002600.

Distler, M. G., and A. A. Palmer, 2012 Role of Glyoxalase 1 (Glo1) and methylglyoxal (MG) in behavior: recent advances and mechanistic insights. Front Genet 3: 250.

Dobin, A., C. A. Davis, F. Schlesinger, J. Drenkow, C. Zaleski et al., 2013 STAR: ultrafast universal RNA-seq aligner. Bioinformatics 29: 15–21.

Eisenberg, E., and E. Y. Levanon, 2013 Human housekeeping genes, revisited. Trends Genet 29: 569–574.

Fu, J., J. J. Keurentjes, H. Bouwmeester, T. America, F. W. Verstappen et al., 2009 System-wide molecular evidence for phenotypic buffering in Arabidopsis. Nat Genet 41: 166–167.

Geisert, E. E., L. Lu, N. E. Freeman-Anderson, J. P. Templeton, M. Nassr et al., 2009 Gene expression in the mouse eye: an online resource for genetics using 103 strains of mice. Mol Vis 15: 1730–1763.

Ghazalpour, A., B. Bennett, V. A. Petyuk, L. Orozco, R. Hagopian et al., 2011 Comparative analysis of proteome and transcriptome variation in mouse. PLoS Genet 7: e1001393.

Haider, S., B. Ballester, D. Smedley, J. Zhang, P. Rice et al., 2009 BioMart Central Portal-unified access to biological data. Nucleic Acids Res 37: W23–27.

Haley, C. S., S. A. Knott and J. M. Elsen, 1994 Mapping quantitative trait loci in crosses between outbred lines using least squares. Genetics 136: 1195–1207.

Hasin-Brumshtein, Y., F. Hormozdiari, L. Martin, A. Van Nas, E. Eskin et al., 2014 Allele-specific expression and eQTL analysis in mouse adipose tissue. BMC Genomics 15: 471.

Hilgers, V., M. W. Perry, D. Hendrix, A. Stark, M. Levine et al., 2011 Neural-specific elongation of 3' UTRs during Drosophila development. Proc Natl Acad Sci U S A 108: 15864–15869.

Huang DA, W., B. T. Sherman and R. A. Lempicki, 2009 Systematic and integrative analysis of large gene lists using DAVID bioinformatics resources. Nat Protoc 4: 44–57.

Keane, T. M., L. Goodstadt, P. Danecek, M. A. White, K. Wong et al., 2011 Mouse genomic variation and its effect on phenotypes and gene regulation. Nature 477: 289–294.

Keen, J. C., and H. M. Moore, 2015 The Genotype-Tissue Expression (GTEx) Project: Linking Clinical Data with Molecular Analysis to Advance Personalized Medicine. J Pers Med 5: 22–29.

Kendig, E. L., Y. Chen, M. Krishan, E. Johansson, S. N. Schneider et al., 2011 Lipid metabolism and body composition in Gclm(-/-) mice. Toxicol Appl Pharmacol 257: 338–348.

Kent, W. J., 2002 BLAT-the BLAST-like alignment tool. Genome Res 12: 656–664.

Lagarrigue, S., L. Martin, F. Hormozdiari, P. F. Roux, C. Pan et al., 2013 Analysis of allele-specific expression in mouse liver by RNA-Seq: a comparison with Cis-eQTL identified using genetic linkage. Genetics 195: 1157–1166.

Lappalainen, T., M. Sammeth, M. R. Friedlander, P. A. T Hoen, J. Monlong et al., 2013 Transcriptome and genome sequencing uncovers functional variation in humans. Nature 501: 506–511.

Lareau, L. F., A. N. Brooks, D. A. Soergel, Q. Meng and S. E. Brenner, 2007 The coupling of alternative splicing and nonsense-mediated mRNA decay. Adv Exp Med Biol 623: 190–211.

Li, H., and R. Durbin, 2010 Fast and accurate long-read alignment with Burrows-Wheeler transform. Bioinformatics 26: 589–595.

Li, H., B. Handsaker, A. Wysoker, T. Fennell, J. Ruan et al., 2009 The Sequence Alignment/Map format and SAMtools. Bioinformatics 25: 2078–2079.

Li, Z., M. K. Mulligan, X. Wang, M. F. Miles, L. Lu et al., 2010 A transposon in Comt generates mRNA variants and causes widespread expression and behavioral differences among mice. PLoS One 5: e12181.

Liang, H., A. R. Cavalcanti and L. F. Landweber, 2005 Conservation of tandem stop codons in yeasts. Genome Biol 6: R31.

Lin, S., Z. Yang, H. Liu and Z. Cai, 2011 Metabolomic analysis of liver and skeletal muscle tissues in C57BL/6J and DBA/2J mice exposed to 2,3,7,8-tetrachlorodibenzo-p-dioxin. Mol Biosyst 7: 1956–1965.

Lonsdale, J., J. Thomas and M. Salvatore, 2013 The Genotype-Tissue Expression (GTEx) project. Nat Genet 45: 580–585.

Lu, A., X. Wangpu, D. Han, H. Feng, J. Zhao et al., 2012 TXNDC9 expression in colorectal cancer cells and its influence on colorectal cancer prognosis. Cancer Invest 30: 721–726.

Maier, T., M. Guell and L. Serrano, 2009 Correlation of mRNA and protein in complex biological samples. FEBS Lett 583: 3966–3973.

Manolio, T. A., F. S. Collins, N. J. Cox, D. B. Goldstein, L. A. Hindorff et al., 2009 Finding the missing heritability of complex diseases. Nature 461: 747–753.

Masel, J., and M. L. Siegal, 2009 Robustness: mechanisms and consequences. Trends Genet 25: 395–403.

Maurano, M. T., R. Humbert, E. Rynes, R. E. Thurman, E. Haugen et al., 2012 Systematic localization of common disease-associated variation in regulatory DNA. Science 337: 1190–1195.

Mckenna, A., M. Hanna, E. Banks, A. Sivachenko, K. Cibulskis et al., 2010 The Genome Analysis Toolkit: a MapReduce framework for analyzing next-generation DNA sequencing data. Genome Res 20: 1297–1303.

Mcmanus, C. J., J. D. Coolon, M. O. Duff, J. Eipper-Mains, B. R. Graveley et al., 2010 Regulatory divergence in Drosophila revealed by mRNA-seq. Genome Res 20: 816–825.

Mitterberger, M. C., G. Kim, U. Rostek, R. L. Levine and W. Zwerschke, 2012 Carbonic anhydrase III regulates peroxisome proliferator-activated receptor-gamma2. Exp Cell Res 318: 877–886.

Miura, P., S. Shenker, C. Andreu-Agullo, J. O. Westholm and E. C. Lai, 2013 Widespread and extensive lengthening of 3' UTRs in the mammalian brain. Genome Res 23: 812–825.

Mozhui, K., L. Lu, W. E. Armstrong and R. W. Williams, 2012 Sex-specific modulation of gene expression networks in murine hypothalamus. Front Neurosci 6: 63.

Pandey, A. K., and R. W. Williams, 2014 Genetics of gene expression in CNS. Int Rev Neurobiol 116: 195–231.

Peirce, J. L., E. J. Chesler, R. W. Williams and L. Lu, 2003 Genetic architecture of the mouse hippocampus: identification of gene loci with selective regional effects. Genes Brain Behav 2: 238–252.

Rozowsky, J., A. Abyzov, J. Wang, P. Alves, D. Raha et al., 2011 AlleleSeq: analysis of allele-specific expression and binding in a network framework. Mol Syst Biol 7: 522.

Satya, R. V., N. Zavaljevski and J. Reifman, 2012 A new strategy to reduce allelic bias in RNA-Seq readmapping. Nucleic Acids Res 40: e127.

Schadt, E. E., S. A. Monks, T. A. Drake, A. J. Lusis, N. Che et al., 2003 Genetics of gene expression surveyed in maize, mouse and man. Nature 422: 297–302.

Schwanhausser, B., D. Busse, N. Li, G. Dittmar, J. Schuchhardt et al., 2011 Global quantification of mammalian gene expression control. Nature 473: 337–342.

She, X., C. A. Rohl, J. C. Castle, A. V. Kulkarni, J. M. Johnson et al., 2009 Definition, conservation and epigenetics of housekeeping and tissue-enriched genes. BMC Genomics 10: 269.

Simpson, J. T., R. E. Mcintyre, D. J. Adams and R. Durbin, 2010 Copy number variant detection in inbred strains from short read sequence data. Bioinformatics 26: 565–567.

Skelly, D. A., G. E. Merrihew, M. Riffle, C. F. Connelly, E. O. Kerr et al., 2013 Integrative phenomics reveals insight into the structure of phenotypic diversity in budding yeast. Genome Res 23: 1496–1504.

Stamatoyannopoulos, J. A., M. Snyder, R. Hardison, B. Ren, T. Gingeras et al., 2012 An encyclopedia of mouse DNA elements (Mouse ENCODE). Genome Biol 13: 418.

Szabo, P. E., and J. R. Mann, 1995 Allele-specific expression and total expression levels of imprinted genes during early mouse development: implications for imprinting mechanisms. Genes Dev 9: 3097–3108.

Tarazona, S., F. Garcia-Alcalde, J. Dopazo, A. Ferrer and A. Conesa, 2011 Differential expression in RNA-seq: a matter of depth. Genome Res 21: 2213–2223.

Vilella, A. J., J. Severin, A. Ureta-Vidal, L. Heng, R. Durbin et al., 2009 EnsemblCompara GeneTrees: Complete, duplication-aware phylogenetic trees in vertebrates. Genome Res 19: 327–335.

Waddington, C. H., 1942 Canalization of development and the inheritance of acquired characters. Nature 150: 563.

Wang, X., R. Agarwala, J. Capra, Z. Chen, D. Church et al., 2010 High-throughput sequencing of the DBA/2J mouse genome. BMC Bioinformatics 11: O7.

Wang, X., K. Mozhui, Z. Li, M. K. Mulligan, F. J. Ingels et al., 2012 A promoter polymorphism in the Per3 gene is associated with alcohol and stress response. Translational Psychiatry 2: 81–85.

Wang, X., Q. Sun, S. D. Mcgrath, E. R. Mardis, P. D. Soloway et al., 2008 Transcriptome-wide identification of novel imprinted genes in neonatal mouse brain. PLoS One 3: e3839.

Ward, L. D., and M. Kellis, 2012 Interpreting noncoding genetic variation in complex traits and human disease. Nat Biotechnol 30: 1095–1106.

Weng, M. P., and B. Y. Liao, 2010 MamPhEA: a web tool for mammalian phenotype enrichment analysis. Bioinformatics 26: 2212–2213.

Williams, E. G., L. Mouchiroud, M. Frochaux, A. Pandey, P. A. Andreux et al., 2014 An evolutionarily conserved role for the aryl hydrocarbon receptor in the regulation of movement. PLoS Genet 10: e1004673.

Wu, Y., E. G. Williams, S. Dubuis, A. Mottis, V. Jovaisaite et al., 2014 Multilayered genetic and omics dissection of mitochondrial activity in a mouse reference population. Cell 158: 1415–1430.

Zhang, L., and W. H. Li, 2004 Mammalian housekeeping genes evolve more slowly than tissue-specific genes. Mol Biol Evol 21: 236–239.

Zhang, Z., D. Xin, P. Wang, L. Zhou, L. Hu et al., 2009 Noisy splicing, more than expression regulation, explains why some exons are subject to nonsense-mediated mRNA decay. BMC Biol 7: 23.

